# A chemical inducer of ribophagy limits the toxicity of ALS-related arginine-rich peptides

**DOI:** 10.64898/2026.03.15.711847

**Authors:** Bartlomiej Porebski, Vanesa Lafarga, Catherine Hansel, Martin Haraldsson, Anabel Saez, Angela Sanchez-Molleda, Saioa Moragon, Maria Haggblad, Louise Lidemalm, Sara Rodrigo, Jordi Carreras-Puijgvert, Iván Ventoso, Miguel Lafarga, Daniela Hühn, Oscar Fernández-Capetillo

**Affiliations:** Division of Genome Biology, Science for Life Laboratory, Department of Medical Biochemistry and Biophysics, Karolinska Institute, Stockholm, Sweden; Chemical Biology Consortium Sweden, Science for Life Laboratory, Department of Medical Biochemistry and Biophysics, Karolinska Institute, Stockholm, Sweden; Genomic Instability Group, Spanish National Cancer Research Centre (CNIO), Madrid, Spain; Novo Nordisk Research Centre, Oxford, UK; Miaker Developments SL, Miaker Bioassays, San Sebastian, Spain; Department of Neurosciences, Biogipuzkoa Health Research Institute, San Sebastian, Spain; Department of Pharmaceutical Biosciences, Uppsala University, Uppsala, Sweden; Pixl Bio AB, Uppsala, Sweden; Centro de Biologia Molecular Severo Ochoa (CSIC-UAM), Departamento de Biologia Molecular, Universidad Autonoma de Madrid (UAM), Madrid, Spain; Department of Anatomy and Cell Biology, University of Cantabria, Santander, Spain; Networking Research Center on Neurodegenerative Diseases (CIBER-NED), Instituto de Salud Carlos III, Madrid, Spain

**Author notes:** **Correspondence**: Oscar Fernandez-Capetillo.

**Keywords:** Chemical screen, ALS, arginine-rich peptides, ribophagy, rRNA

## Abstract

*C9ORF72* intronic repeat expansions are the most frequent mutation found in Amyotrophic Lateral Sclerosis (ALS), producing toxic arginine-rich dipeptides (DPR) that disrupt RNA metabolism and trigger the accumulation of orphan ribosomal proteins (RP). Through a large phenotypic chemical screen, we identified “SALSa”, a novel compound that mitigates DPR toxicity. Mechanistically, SALSa acts as a chemical inducer of ribophagy, a specialized form of autophagy that promotes RP clearance. Interestingly, this effect is unrelated to mTOR inhibition, the main regulator of autophagy. In contrast, this is due to an effect of the drug in ribosome biogenesis, which triggers a protective response to clear defective ribosomes. Accordingly, SALSa accumulates in nucleoli and perturbs the final steps of rRNA maturation. SALSa reduces DPR toxicity in differentiated neurons and significantly extends lifespan in a *Drosophila melanogaster* model of *C9ORF72* ALS. These findings suggest that stimulating ribophagy could be beneficial for pathologies associated to dysfunctional ribosome biogenesis, including *C9ORF72* ALS.

## Introduction

Amyotrophic lateral sclerosis (ALS) is a fatal neurodegenerative disease driven by the progressive loss of motorneurons (MN)^1^. Current approved therapies include Riluzole, Edaravone and the antisense oligonucleotide Tofersen for patients with mutations in SOD1^2^. Unfortunately, and despite recent promising data for Tofersen-treated ALS patients^3^, particularly for those that start the treatment early, disease prognosis remains very poor. In addition, and despite its low incidence of ∼2 per 100,000 person-years worldwide^4^, the number of affected patients is rapidly increasing mainly due to the ageing of our populations^5^. Thus, additional therapies that explore novel pathways or targets are urgently needed.

Part of the complexity in ALS lies in its multifactorial origin, as the disease has been associated to mutations in more than 30 genes and hundreds of genetic variants^6^. In addition, only 10% of ALS cases are hereditary, while the rest are thought to be the result of environmental exposures or unknown genetic determinants. Nevertheless, mutations in *C9ORF72* constitute the main hereditary cause of ALS, being present in up to 40% of the familiar cases and 5-10% of the sporadic ones^7,8^. The mutation is an expansion of a GGGGCC hexanucleotide repeat (HRE) in the first intron of *C9ORF72*, and several independent models have been proposed to explain the pathogenicity of the mutation^9^. These include the loss of C9ORF72 function and the toxicity of HRE-containing RNAs. In addition, HRE sequences are translated by repeat-associated non-AUG (RAN) translation, giving rise to five dipeptide repeats (DPRs)^10,11^. Importantly, arginine-rich DPRs ((PR)n and (GR)n)) accumulate at nucleoli and are toxic for eukaryotic cells, providing a straight-forward model to explain the mechanism of the *C9ORF72* mutation^12^.

In recent years, we have tried to contribute to deciphering the mechanism by which arginine-rich DPRs cause cellular toxicity. Our initial work revealed that the very high affinity of these peptides for RNA leads to a collective displacement of RNA-binding proteins from all cellular RNAs^13^. Subsequently, we found that, by sequestering rRNA and limiting its maturation, (PR)n peptides impair the assembly of novel ribosomal particles, leading to a generalized accumulation of orphan ribosomal proteins (oRPs)^14^. Due to their abundance, oRP accumulation is particularly toxic for eukaryotic cells, as it saturates proteostasis and aggregate clearance machineries^15^. However, therapies that target this phenomenon are missing. In this regard, in 2019 we reported an initial drug-repurposing screen which allowed us to identify several drugs capable of reducing (PR)n toxicity in cells and developing zebrafish^16^. Building up on this approach, we here present the results of a much larger chemical screening campaign oriented to find novel chemicals capable of reducing the toxicity of arginine-rich DPRs.

## Results

### A chemical screen to identify modulators of DPR toxicity

To identify drugs capable of alleviating DPR toxicity, we used a previously described U2OS cell line (U2OS^PR97^)^16^. In these cells, treatment with doxycycline (dox) triggered a widespread expression of (PR)_97_ peptides which, as reported, accumulated at nucleoli (**Fig. 1A**) and led to a significant loss of cell viability (**Fig. 1B**). Using this model, we conducted a chemical screen of 35,211 uncharacterized compounds (see methods for library composition). To do so, U2OS^PR97^ cells were seeded in 384-well plates, and exposed to 10 μM of the compounds for 72h, at which point cell viability was measured using CellTiter-Glo (**Fig. 1C**). Cell viability dropped to around 40% in control wells upon dox addition, and the screening was technically robust as measured by its Ź (0.55) or a low coefficient of variation (**Fig. S1A-C**). 51 compounds were selected from this screen on the basis of increasing viability over 3 standard deviations (SD) from control wells and having a Z > 2 (based on all compounds) (**Fig. 1D**).

**Figure 1.**
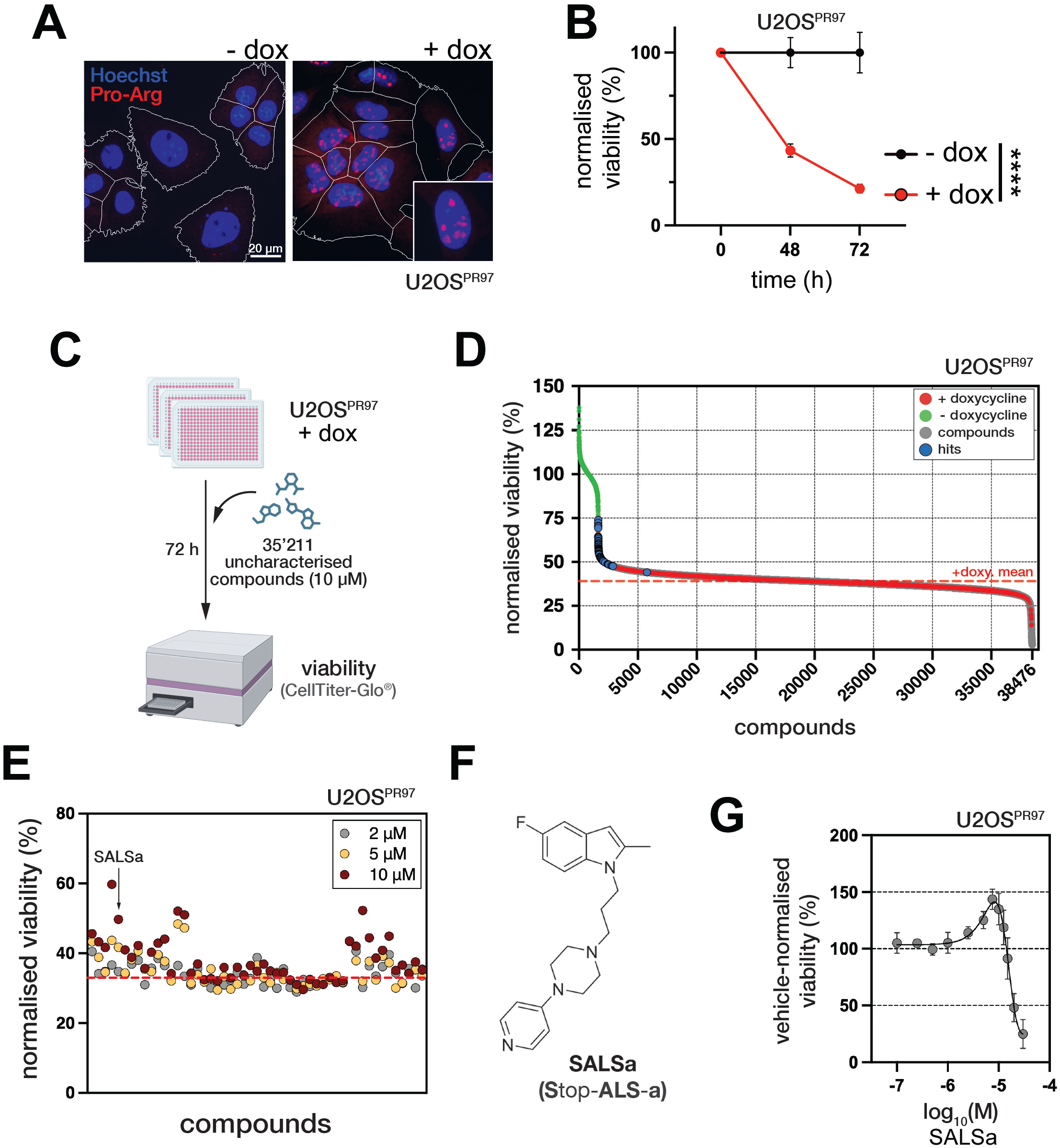
A chemical screen for modulators of DPR toxicity. A. Microscopy images of U2OS^PR97^ cells untreated (left) or treated with 20 ng/ml dox for 24 hours (right). Cells were stained with Hoechst 3342 (blue) to visualize nuclei, and anti-Pro-Arg antibody (red). White outlines delineate cell borders. Scale bars, 20 *µ*M. An inset is shown to illustrate the nucleolar accumulation of (PR)_97_ peptides. B. Time-course illustrating the dox-dependent decrease of viability in U2OS^PR97^ cells. Viability was assessed by counting nuclei stained with Hoechst 3342. Data were normalised to the uninduced sample within each time-point. Plotted values represent mean from 3 replicates. Statistical significance was determined by two-way ANOVA. **** p < 0.0001 C. Pipeline of the chemical screen. U2OS^PR97^ cells were seeded in the presence of 20 ng/ml dox onto 384-well plates pre-spotted with a chemical library consisting of 35,211 uncharacterised drug-like small molecule compounds. After 72 hours, viability was measured using Cell-TiterGlo®. D. Representation of the screening dataset. Green dots represent positive controls (without dox and vehicle-treated); red dots represent negative controls (i.e. dox-treated and vehicle-treated); grey dots represent tested compounds and orange dots represent hits (51 compounds). E. Validation of primary screen hits in a 3-point dose response experiment. Plotted is the viability normalised against dox-treated U2OS^PR97^ cells (N = 3). Red dashed line indicates viability of the negative control. F. Structure of our top hit compound, SALSa. G. Dose-response curve of SALSa in dox-treated U2OS^PR97^ cells. Plotted is the viability normalised against dox-treated cells (N = 3).

Hits from the primary screen were subsequently tested in a validation screen at 3-doses using the same assay, bringing down the list to 10 compounds that consistently increased the viability of dox-treated U2OS^PR97^ cells (**Fig. 1E**). Of note, chemical screens based on dox-inducible expression systems can yield false-positives due to compounds that prevent the expression of the reporter^17^. Thus, in another validation, we discarded compounds that reduced dox-induced (PR)_97_ levels in U2OS^PR97^ cells, or that decreased EGFP levels in an independent U2OS^EGFP^ cell line harboring dox-induced EGFP (**Fig. S1D**). These analyses led us to focus on one compound, which we named “SALSa” (<u>S</u>top <u>ALS</u> <u>a</u>), as our top candidate (**Fig. 1F**). SALSa treatment led to a dose-dependent increase of viability in dox-treated U2OS^PR97^ cells, yet it was toxic at higher doses (**Fig. 1G**).

Structure-activity relationship (SAR) identified that the indole core ring of the molecule tolerated small substitutions without losing potency, which allowed us to generate derivatives for imaging and target identification (**Fig. S2A**). In contrast, the two-ring system was more sensitive to changes, requiring a basic composition at the distal end of the aryl ring (**Fig. S2B**). We also identified substitutions in SALSa that increased its potency, yet all faced toxicity at higher doses (**Fig. S2C**). In light of our mechanistic experiments (see below), our interpretation of this biphasic behavior is that the compound is likely affecting an essential target, in such a way that a partial targeting might be beneficial, but its complete inactivation is deleterious. Together, these experiments allowed us to identify SALSa, as a novel chemical entity that limits the toxicity of arginine-rich peptides.

### SALSa has a distinct effect on ribosome metabolism

We then tried to decipher the mechanism of action of SALSa. First, we conducted RNAseq in cells treated with SALSa at 10 μM for 6h. Analysis of the SALSa-induced transcriptome showed a distinct upregulation of ribosomal genes (**Fig. 2A**). Consistently, “Ribosome” was the top upregulated pathway that emerged from enrichment analyses ran against the “Kyoto Encyclopedia of Genes and Genomes” (KEGG) (**Fig. 2B**).

**Figure 2.**
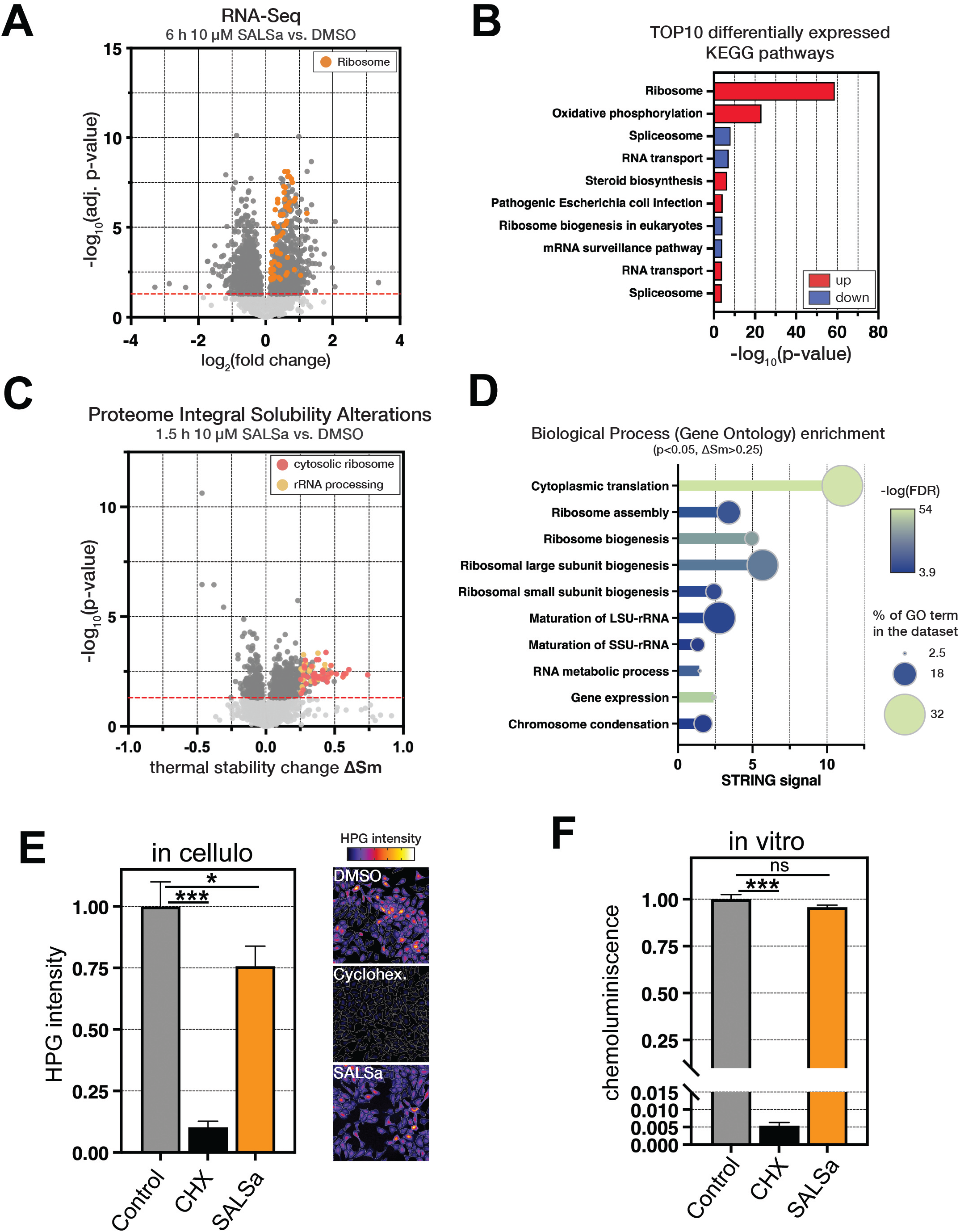
SALSa modulates translation and ribosome biogenesis. A. Volcano plot representing gene expression changes in MCF-7 cells treated with SALSa at 10 *µ*M for 6 h. Horizontal red line marks the adjusted *p*-value threshold. Dark grey dots represent significantly changed genes and orange dots represent ribosomal genes. B. Enrichment analysis of Kyoto Encyclopedia of Genes and Genomes (KEGG) pathways upregulated (red) or downregulated (blue) in SALSa-treated cells. C. Volcano plot showing proteome solubility changes in U2OS cells treated with SALSa at 10 *µ*M for 1.5 h, assessed with the Proteome Integral Solubility Alterations (PISA) assay. Horizontal red line marks the *p*-value threshold. Coloured dots represent the most enriched clusters within significantly affected proteins. D. Enrichment analysis of Gene Ontology Biological Processes among the proteins most significantly affected by SALSa. STRING analysis using proteins with p<0.05 and ΔSm>0.25 from the PISA assay. E. *In cellulo* translation rate quantification using L-homopropargylglycine (HPG) incorporation assay in U2OS cells treated for 6 h with DMSO, 100 ng/ml Cycloheximide (CHX) or 10 *µ*M SALSa. Plotted values are mean of 3 replicates normalised to the DMSO sample. Statistical significance was determined by two-way ANOVA. *, p < 0.05; ***, p < 0.001 F. *In vitro* translation rates by measuring the translation of a luciferase mRNA in rabbit reticulocyte lysates in the presence of DMSO, 100 ng/ml CHX or 10 *µ*M SALSa. Plotted values are mean of 3 replicates normalised to the DMSO sample. Statistical significance was determined by two-way ANOVA; ns, non-significant; ***, p < 0.001.

Next, to get a more direct view of the proteins that are directly affected by SALSa, we conducted a proteome integral solubility alterations (PISA) assay^18^, which measures how a drug affects the thermal stability of proteins. PISA was conducted in U2OS cells treated with 10 μM SALSa for 1.5h. This analysis revealed an enrichment of cytosolic RPs as well as factors involved in rRNA processing (**Fig. 2C**). Along these lines, enrichment analysis of the Gene Ontology “Biological Process” category using the proteins with a significant alteration of their thermal stability (τιSm > 0.25; p < 0.05), identified “cytoplasmic translation” as well as pathways related to rRNA maturation and ribosome biogenesis as those most affected by SALSa (**Fig. 2D**).

Consistent with transcriptomic and proteomic data, SALSa reduced translation levels in U2OS cells, as measured by the incorporation of L-homopropargylglycine (HPG) (**Fig. 2E**). However, the compound did not reduce translation rates in vitro, arguing against a direct inhibitory effect on the translation machinery (**Fig. 2F**). Sucrose gradient fractionations also failed to detect an effect of SALSa on polysome assembly (**Fig. S3A**), and the compound did not impact on the two main signaling pathways that regulate translation; mTOR and the integrated stress response (ISR) (**Fig. S3B, C**). Together, these data suggest that the influence of SALSa on translation is not direct, but rather the consequence of an effect of the chemical in rRNA processing and/or ribosome biogenesis.

### SALSa is a chemical inducer of ribophagy

Given that omics data indicated an impact of SALSa on ribosomes, we evaluated whether the drug affected RP localization. Immunofluorescence (IF) microscopy revealed that the compound induced the formation of cytoplasmic RPL11 and RPL37A foci, which were reminiscent of the localization of lysosomes or autophagic vesicles. Indeed, RPL11 and RPL37A foci colocalized with Lysosomal-Associated Membrane Protein 1 (LAMP1) (**Fig. 3A**). Transmission electron microscopy (TEM) confirmed the formation of autophagic vesicles filled with RPs, indicating that SALSa induced ribophagy (**Fig. 3B, S4A**). To further confirm the activation of ribophagy, we used a previously described reporter system based on the expression of RPS3 harboring a C-terminal pH-sensitive Keima tag (RPS3^Keima^)^19^. As previously reported, mTOR inhibition led to the lysosomal accumulation of RPS3Keima, which was similarly observed upon treatment with SALSa (**Fig. 3C, D**).

**Figure 3.**
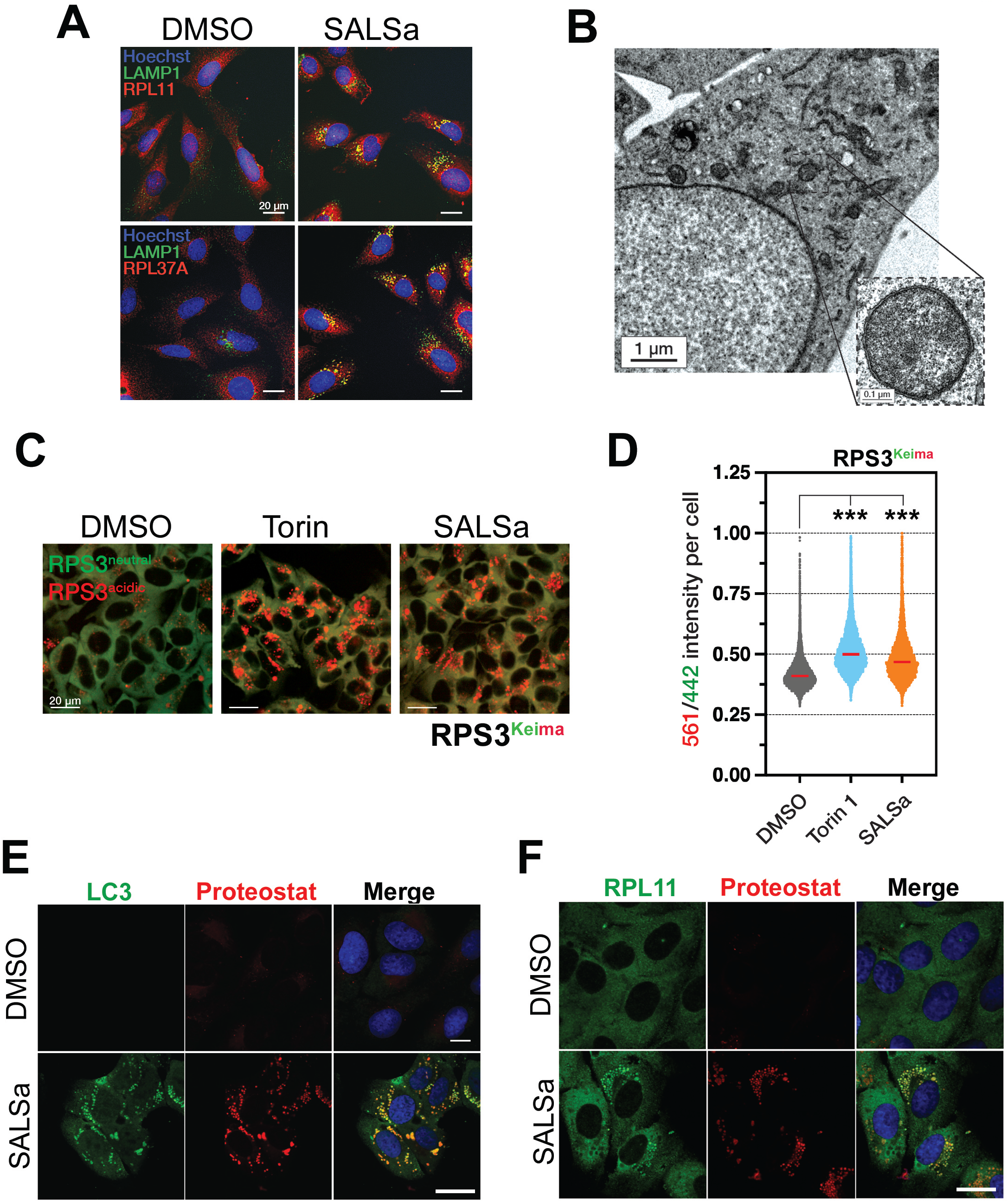
SALSa induces ribophagy. A. Microscopy images of U2OS cells treated with DMSO or with 10 *µ*M SALSa for 6 h and stained with Hoechst 334a (blue) to visualise nuclei, anti-LAMP1 antibody (green) to visualise lysosomes, and anti-RPL11/RPL37A antibodies (red) to visualise ribosomes. Scale bar, 20 *µ*m. B. Transmission electron microscopy image of U2OS cells treated with 10 *µ*M SALSa for 4 h. Inset shows magnification of autophagosomes filled with ribosomes, indicating ongoing ribophagy. C. Microscopy images of U2OS-RPS3^Keima^ cells treated for 48 h with DMSO, 250 nM Torin1 or 10 *µ*M SALSa. Green signal represents cytoplasmic ribosomes (neutral pH), while red signal represents ribosomes undergoing autophagic clearance (acidic pH). Scale bar, 20 *µ*m. D. Quantification of C. For each cell, ratio between ‘acidic’ and ‘neutral’ Keima signal is plotted. Statistical significance was determined by two-way ANOVA. ***, p < 0.001. E. Microscopy images of U2OS cells treated for 24 h with DMSO or 10 *µ*M SALSa and stained with Hoechst 3342 (blue) to visualise nuclei, anti-LC3 antibodies (green) to mark autophagosomes and proteostat (red) to detect protein aggregates. Scale bar, 10 *µ*m. F. Microscopy images of U2OS cells treated for 24 h with DMSO or 10 *µ*M SALSa and stained with Hoechst 3342 (blue) to visualise nuclei, anti-RPL11 antibodies (green) and proteostat (red) to detect protein aggregates. Scale bar, 10 *µ*m.

Interestingly, use of a dye that stains protein aggregates (proteostat) revealed that SALSa induced the accumulation or protein aggregates containing RPs, that often localized at autophagic vesicles (**Fig. 3E, F**). Furthermore, TEM analyses identified RP-aggregates and ribophagy vesicles in SALSa treated cells (**Fig. S4A**). Next, we used a cell line expressing Halo-tagged RPL29, which enables tracking ribosomal subpopulations by sequential pulse labelling with two different fluorophores (TMR and 11R)^20^. These experiments revealed that SALSa specifically affected the levels of RPs produced in the presence of the drug, but not of those that were already present before the treatment (**Fig. S4B, C**). Together, these data suggest that SALSa affects ribosome biogenesis in a manner that triggers ribophagy to promote the clearance of dysfunctional ribosomes.

### SALSa limits the final steps of rRNA maturation

Based on our SAR studies, we generated an alkyne-derivative of SALSa (SALSa^alk^) suitable for Click-chemistry (**Fig 4A**). The compound retained its effects on limiting (PR)_97_ toxicity (**Fig. 4B**). IF experiments revealed that SALSa^alk^ accumulated in nucleoli (stained by NPM1) and lysosomes (stained by LAMP1) (**Fig. 4C**). Despite its presence in nucleoli, the compound did not have a major effect on nucleolar size or integrity (**Fig. S5A**), or on rDNA transcription (**Fig. S5B**). Moreover, and in contrast to the RNA polymerase I inhibitor Actinomycin D, the compound failed to activate the nucleolar stress response as measured by p53 expression (**Fig. S5C**).

**Figure 4.**
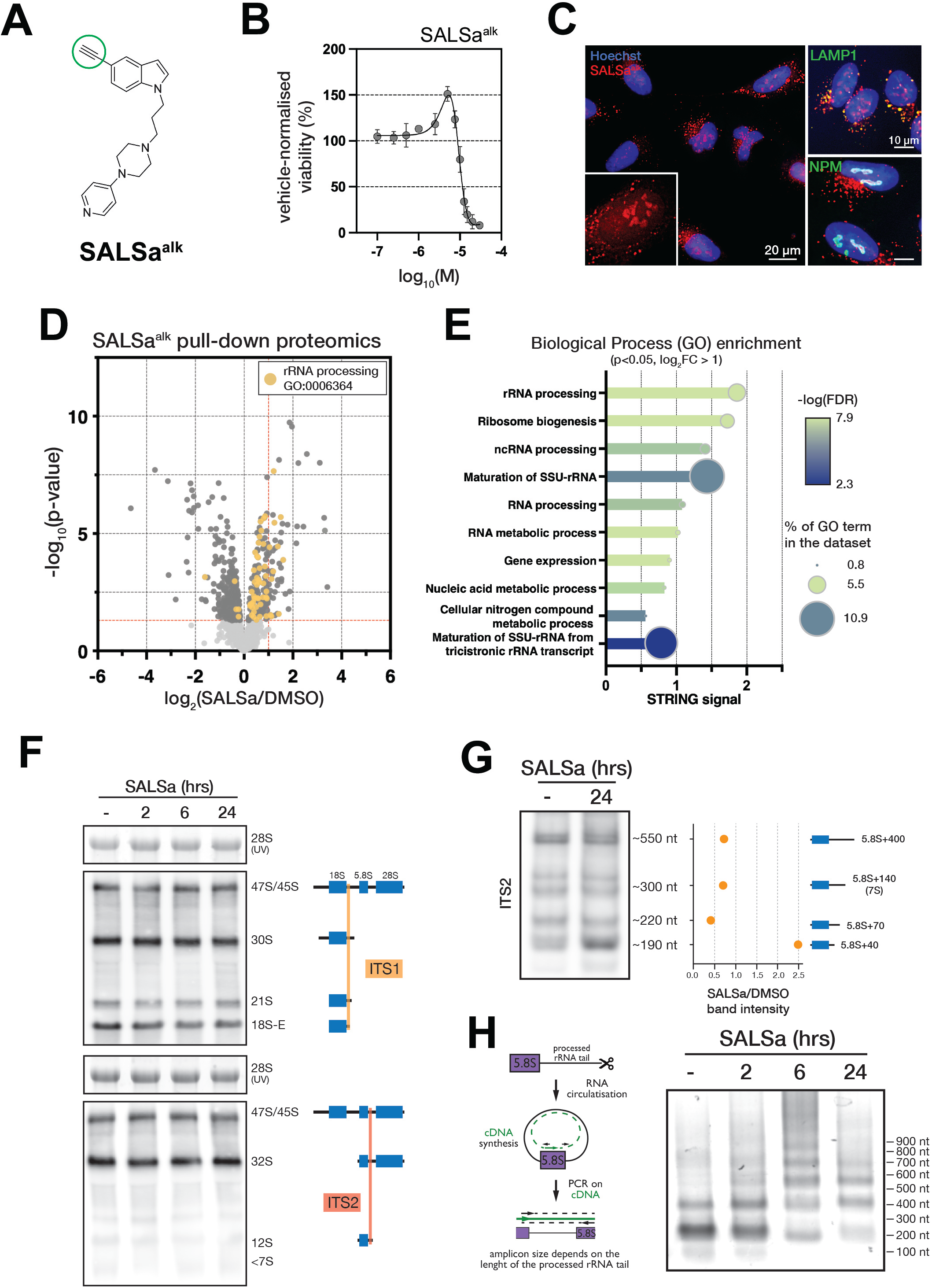
SALSa accumulates at nucleoli and impairs 5.8S rRNA maturation. A. Structure of SALSa functionalised with an alkyne group (SALSa^alk^). B. Activity of SALSa^alk^ in rescuing toxicity in U2OS^PR97^ cells. Plotted is the viability normalised against dox-treated cells (N = 3). C. Microscopy images of U2OS cells treated with 10 *µ*M SALSa^alk^ for 2 h. Cells were stained with Hoechst 3342 (blue) to visualise nuclei, SALSa^alk^ was visualised by Click chemistry coupling of Alexa^647^ fluorophore (red). Counter-staining with antibodies (green) was done to visualise lysosomes (LAMP1) or nucleoli (NPM). Scale bar as indicated. D. Volcano plot of the proteomic analysis of SALSa^alk^ interacting proteins. Horizontal red line indicates the p-value threshold (<0.05), while the vertical red line marks a 2x fold-change. Yellow dots represent proteins in the data set belonging to the ‘rRNA processing’ Gene Ontology Biological Process term, the most significantly enriched in the data set. E. Enrichment analysis of Gene Ontology Biological Processes among the proteins enriched at least 2-fold in the SALSa^alk^ pull-down sample compared to the mock pull-down. The analysis was done using STRING. F. Northern blotting of rRNA intermediates in U2OS cells treated for 24 h with 10 *µ*M SALSa. Total cellular RNA was resolved using agarose gel electrophoresis, blotted onto nitrocellulose membrane, and biotinylated probes directed to Internal Transcribed Spacers (ITS1 and ITS2) in rRNA primary transcript (see schematic) were hybridized. The image of GelRed-stained 28S mature rRNA is provided as a loading control. G. As in F, but total cellular RNA was resolved on a polyacrylamide gel to enable the analysis of 5.8S rRNA maturation intermediates. H. Analysis of 5.8S rRNA maturation in U2OS cells treated for 24 h with 10 *µ*M SALSa using a circularised RNA RT-PCR assay. An scheme illustrating the assay is provided to the left of the data. Total RNA was circularised with T4 RNA ligase, followed by reverse transcription with an internal 5.8S primer and end-point PCR on the resulting cDNA. Reactions were resolved by agarose gel electrophoresis.

We then used SALSa^alk^ for chemical-affinity purification followed by mass spectrometry (MS). Proteomic analyses of SALSa^alk^ interactors identified “rRNA processing” as the most significantly enriched pathway (**Fig. 4D, E**). Northern blotting of rRNA intermediates failed to detect major changes in rRNA processing induced by SALSa (**Fig. 4F**). However, running these analyses on polyacrylamide gels to analyze 5.8S maturation revealed an accumulation of unprocessed intermediates (**Fig. 4G**). Consistent with this, a quantitative rRNA maturation assay based on RNA circularization revealed that SALSa hindered the exonucleolytic processing of the Internal Transcribed Spacer 2 (ITS2) tail from the 5.8S rRNA (**Fig. 4H**). These experiments show that SALSa accumulates in nucleoli, where it impairs the final steps of rRNA maturation.

### Impact of SALSa in human neurons and fly ALS models

Finally, we evaluated the effects of SALSa in additional cellular and animal models. First, and consistent with our findings in U2OS cells, SALSa reduced (PR)_97_ toxicity in neuronally-differentiated human SH-SY5Y cells (SH-SY5Y^PR97^) (**Fig. 5A, B**). Of note, the dose required to rescue differentiated neurons was in the low nanomolar range. Beyond in vitro data, and given that the *C9ORF72* mutation has been proposed to drive motor neuron loss for reasons beyond arginine-rich DPRs^21^, we tested SALSa in 2 independent models of *Drosophila melanogaster* with expression of the human *C9ORF72* hexanucleotide repeat expansion either in the central nervous system or in muscles. The mutation compromised survival in both models, which was significantly alleviated upon SALSa treatment (**Fig. 5C, D**). In summary, the experiments presented in this manuscript indicate that by limiting the final steps of rRNA maturation, SALSa induces the clearance of RPs which is protective in the context of pathologies presenting abnormal ribosome biogenesis such as C9ORF72 ALS.

**Figure 5.**
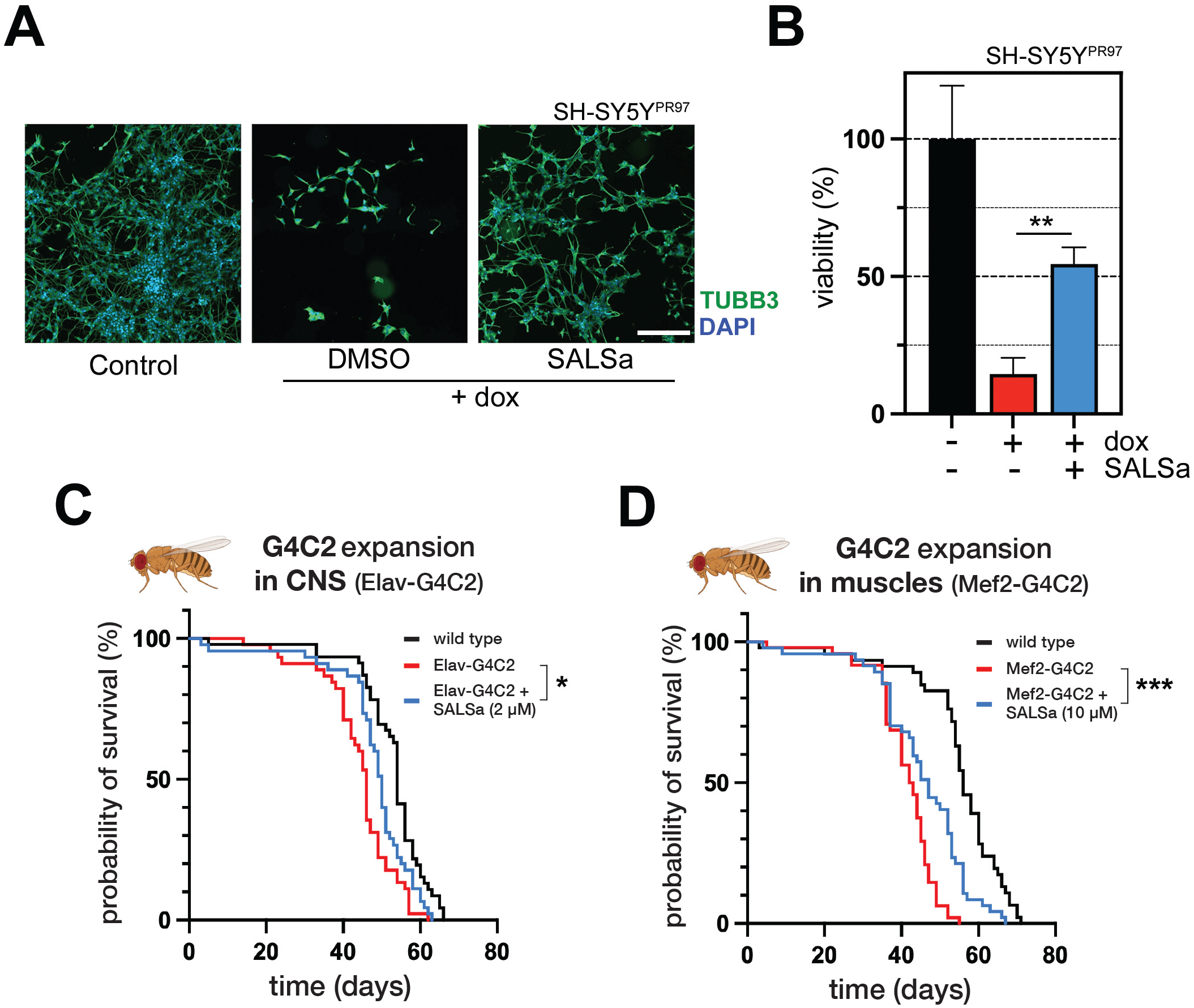
SALSa limits toxicity in human and fly ALS models. A. Microscopy images of dox-inducible SH-SY5Y^PR97^ cells differentiated into neurons and treated with DMSO or 1 nM SALSa. Cells were stained with Hoechst 3342 (blue) to visualise DNA and an anti-βIII Tubulin (TUBB3) antibody to mark cell body (green). B. Effect of SALSa (nM) on the viability of SH-SY5Y^PR97^ cells differentiated into neurons. Plotted values represent the mean of 3 replicates normalised to the uninduced sample. **, p < 0.01. C. Probability of survival of a *Drosophila melanogaster* ALS model expressing the *C9ORF79*-associated G4C2 HRE in the central nervous system. Flies were treated with DMSO (red) or with SALSa (blue). Survival of wild type line is shown in black. Statistical significance was determined by Mantel-Cox test. *, p < 0.05. D. Probability of survival of a *Drosophila melanogaster* ALS model expressing the *C9ORF79*-associated G4C2 HRE in muscle tissue. Flies were treated with DMSO (red) or with SALSa (blue). Survival of wild type line is shown in black. Statistical significance was determined by Mantel-Cox test. ***, p < 0.001.

## Discussion

Despite the abundance of mutations related to RNA binding proteins in ALS or other neurodegenerative diseases^22–24^, how these mutations lead to pathology or how can they be targeted for therapies remains largely unknown. This is best exemplified by the fact that no effective therapies have been yet developed to target TDP-43, an abundant RNA binding protein that forms aggregates in more than 95% of ALS cases^25,26^, becoming the main molecular hallmark of the disease. Multiple of lines of evidence suggest that the pathogenicity driven by the most frequent mutation in ALS, the *C9ORF72* HRE, is also related to RNA biology ^27–32^. In this regard, our works revealed that this is mediated by the high affinity of arginine-rich peptides for RNA^13^ which, by preventing the formation of new ribosomes, trigger a toxic accumulation or orphan ribosomal proteins^14^. The question is now, how to limit the toxicity of this phenomenon.

So far, strategies to target RNA binding proteins have focused on the use of ASOs^33^ and, in the case of *C9ORF72,* in limiting the expression of the HRE-containing RNA and/or in reducing the production of the DPRs^34–42^. Our study here, presents a novel strategy, that is based on the stimulation of selective autophagic pathways that can clear RP aggregates arising from dysfunctional ribosome biogenesis. In this regard, we and others have already reported beneficial effects of inhibiting mTOR, and thereby stimulating autophagy, in reducing DPR toxicity^14,43^. However, mTOR regulates several signaling pathways beyond autophagy, and its inhibition, despite promoting longevity and being neuroprotective in multiple animal models of research, can also have negative effects in the long term^44^. Accordingly, the positive effects of mTOR inhibition in humans are counterbalanced by effects in other aspects such as insulin signaling or immunosuppression. Our identification of SALSa provides a proof-of-concept example that stimulation of selective autophagy pathways, such as ribophagy, offers an alternative approach to promote the clearance of protein aggregates associated to neurodegeneration.

In this context, is important to highlight that SALSa induces ribophagy in an mTOR-independent manner. While the specific target of SALSa remains to be identified, our current working model is that, by accumulating at nucleoli, the compound has a mild effect in limiting the final steps of ribosome biogenesis. By doing so, it activates a stress-protective response which, among others, stimulates ribophagy to facilitate the clearance of dysfunctional ribosomes. The biphasic mode of action of the chemical series, by being toxic at higher doses, suggests that the drug is targeting an essential gene. Noteworthy, recent works have revealed that targeting rRNA processing factors such as fibrillarin (FBL) can have long-term beneficial effects and promote longevity^45^.

The idea around the mild inhibition of an essential target to trigger protective cellular responses with a long-term impact is not new. This is the basis of hormesis, the concept that an early mild stress can protect from disease later in life^46–48^. An interesting example in ALS is provided by the finding that exposure to the neurotoxin beta-methylamino-L-alanine (BMAA) in young mice, delays disease onset and progression in an SOD1 model of the disease^49^. We currently hypothesize that this is the mechanism behind the protective effects of SALSa, yet further work is necessary to elucidate the signaling pathways that link SALSa to ribophagy, and to further characterize the therapeutic potential of this approach in the context of neurodegeneration or other age-related pathologies.

## Materials and Methods

### Cell lines

U2OS (ATCC) and HCT116-RPL29-HaloTag9 (a kind gift from Prof. Wade Harper, Harvard Medical School, USA) cell lines were grown in Dulbecco’s modified Eagle’s medium (Gibco, 31966021) supplemented with 10% foetal bovine serum (FBS; Sigma-Aldrich, F7524) and 1% penicillin-streptomycin (Gibco, 15140122). MCF-7 cells (ATCC) were grown in RPMI 1640 (Gibco, 11875093) supplemented with 10% FBS and 1% penicillin-streptomycin. Inducible U2OS^PR97 16^ and U2OS^EGFP 17^ and grown in DMEM supplemented with 1% penicillin-streptomycin, 10% tetracycline-free FBS (Takara Bio 631106), and 2 ug/ml Blasticidin (InvivoGen, ant-bl-05). Inducible SH SY5Y^PR97^ were generated in the same way as the U2OS^PR97^; the cells were grown in DMEM/F12 (Gibco, 11320033) supplemented with 10% tetracycline-free FBS (PAN Biotech, P30-3602). To generate U2OS-RPS3^Keima^ cells, U2OS cells were infected with pLENTI_RPS3_Keima (Addgene #127140)^50^ and selected with Puromycin for 2 weeks to generate clones.

### Microscopy

To prepare samples for fluorescence microscopy, cells were fixed with 4% formaldehyde for 15 minutes at room temperature, followed by 10-min permeabilization with 0.1% Triton X-100/PBS, 30 min blocking with blocking buffer (3% BSA/0.1% Tween-20/PBS), overnight incubation with primary antibodies at 4°C, and a 1-h incubation with secondary antibody solution supplemented with 2 *µ*M Hoechst 3342 (Thermo Scientific, 62249) and 1 *µ*M CellTracker Orange CMTMR Dye (Thermo Scientific. C2927). PBS washes were done between each step. Quantitative image analysis was performed using CellProfiler ^51^. For the analysis of protein aggregates, cells were fixed with 4% PFA and permeabilized with 0.5% Triton X-100 for 15 min. Cells were then stained with the PROTEOSTAT dye (Enzo, Enz-51023; 1:10,000 dilution) for 30 min. After PROTEOSTAT staining, cells were blocked with 5% PBS–BSA and incubated with the indicated antibodies. All antibodies used in this study are listed in **Table S1**.

### Western blotting

Cells were washed with PBS, scraped on ice into RIPA buffer (Thermo Scientific, PI-89901), sonicated, and centrifuged at 14,000 x g for 15 min at 4°C. Supernatant was recovered and protein concentration was measured using the DC Protein Assay Kit II (Bio-Rad, 5000112). Lysates were boiled in the NuPAGE LDS Sample Buffer (Invitrogen, NP0007) supplemented with NuPAGE Reducing Agent (Invitrogen, NP0009). Samples were electrophoresed on a Bis-Tris gel and electroblotted onto a nitrocellulose membrane (Bio-Rad, 1704270). Membranes were blocked in 5% milk/TBS-Tween buffer and incubated overnight at 4°C with primary antibodies. On the following day, membranes were washed in TBS-Tween, incubated with secondary antibody coupled to horseradish peroxidase for 1 h at room temperature (RT), and washed again in TBS-Tween. Signal was developed using a kit (Thermo Scientific, 10220294) and detected in an Amersham Imager 600 (GE Healthcare). A full list of the antibodies used in this study is available in **Table S1**.

### Screening

For the primary screen, U2OS^PR97^ cells were seeded onto 384-well plates (Corning, 3765) that have been pre-spotted with a screening library of 35,211 small molecules from the Primary Screening Set of the Chemical Biology Consortium Sweden. Final concentration of library compounds was 10 *µ*M; DMSO was used as a negative control. The expression of Pro-Arg dipeptides was induced by 20 ng/ml doxycycline added together with cells. Samples without doxycycline were used as a positive control. Each plate contained 16 wells of both controls. After 72 hours, CellTiter-Glo reagent (Promega, G7571) was added, plates were incubated for 10 minutes, and the luminescence was read using Infinite M200PRO plate reader (Tecan). Data processing was done using KNIME Analytics Platform^52^. Luminescence reading for each compound was normalised to the mean of positive control samples within the same plate. Hit calling criteria were as follows: viability_compound_ > NegC_mean_ + 3 x NegC_stdev_, and Z-score_compound_ > 2 (Z-scores were calculated using the screening library samples only). Potential hits were validated with the same assay setup, but with different doses and a different viability readout (nuclei visualisation using high-content microscopy).

### RNA-Seq

Total RNA was extracted from drug-treated MCF-7 cells using PureLink RNA Mini kit (Invitrogen) according to manufacturer’s protocol. The sequencing library was constructed with the QuantSeq 3’ mRNA-Seq Library Prep Kit (Lexogen), and approximately 10 million reads were obtained by Illumina sequencing. Differential expression and functional analyses were done using BioJupies^53^.

### Proteome Integral Solubility Alterations (PISA) assay

U2OS cells seeded in T25 flasks (3 replicates per condition) were treated for 90 minutes with DMSO or 10 *µ*M SALSa. Samples were processed according to the PISA method^18^. Briefly, cells were detached, washed twice in PBS, and resuspended in PBS with Halt protease inhibitors (ThermoFisher). Cell suspension was aliquoted and a temperature gradient was applied to induce protein precipitation. Samples were then pooled and proteins were extracted by 5 freeze-thaw cycles in liquid nitrogen in the presence of 0.4% NP-40. Soluble proteins were isolated by ultracentrifugation, and the concentration was measured with Micro BCA assay (Thermo Scientific). Samples (50 *µ*g) were reduced, alkylated and precipitated using ice-cold acetone. Protein pellets were sequentially digested using LysC and trypsin. Digested samples were labelled with TMTpro 16-plex reagent kit. The final multiplex peptide sample was cleaned and desalted. Peptides were separated by capillary reversed phase chromatography at high pH and collected to 48 fractions using a capillary UPLC system. Each fraction was speed-vacuum dried, resuspended in 0.1% FA/2% ACN buffer and analyzed by nanoLC-MS/MS, as previously described (Gaetani et al., 2019). Briefly, an Orbitrap Q-Exactive HF mass spectrometer (Thermo Fisher) equipped with an EASY-Spray source and Ultimate 3000 UPLC was employed. Samples were desalted on a PepMap C18 trap column and separated on a 50 cm EASY-Spray C18 column (300 nL/min, 55°C) using a binary solvent system (0.1% FA, 2% ACN in solvent A; 98% ACN, 0.1% FA in solvent B). Elution used a 3–26% B gradient over 97 min and 26–95% over 9 min, followed by a 5 min wash and re-equilibration. MS data were acquired in data-dependent mode with a 45 s dynamic exclusion, full MS scans (m/z 375–1500) at 120000 resolution, followed by HCD MS/MS (33% NCE) on the top precursors (charge 2+–6+) detected at 45000 resolution. MaxQuant Software v1.6 was utilized for the database search and quantification against the Uniprot Homo sapiens (Human) protein database UP000005640. Cysteine carbamidomethylation was set as a fixed modification, along with TMT-related modifications, methionine oxidation, deamidation of arginine and asparagine as variable modifications. Enzyme specificity was defined as trypsin with a maximum of two missed cleavages. A 1% false discovery rate was employed as a filter at both the protein and peptide levels. Contaminants and reversed-hit peptides were removed, and only proteins with at least two unique peptides were included in the quantitative analysis. Proteins with missing values were eliminated. The quantified abundance of each protein in each sample (labeled with a different TMT) was normalized to the total intensity of all proteins in that sample. For each protein in each compound replicate, the normalized protein abundance was divided by the average abundance of that protein in the vehicle-treated replicates. The average ratio across replicates of each compound compared to vehicle control was calculated, and the and τιSm and Log_2_ values of these ratios were determined. Two-tailed Student’s t-test was employed to calculate the p-value, assuming a non-zero ratio.

### Translation assay *in cellulo*

Translation rate in cells was assayed with the use of Click-iT HPG Alexa Fluor Protein Synthesis Assay kit (Invitrogen, C10428) according to manufacturer’s protocol. In short, U2OS cells were seeded onto 96-well plates (Falcon, 353219), allowed to attach overnight, and treated for 6 hours with DMSO, SALSa or cycloheximide. Cells were washed with PBS and incubated 30 minutes with methionine-free DMEM (Gibco, 21013024), followed by 30-minute incubation in the presence of Click-iT Homopropargylglycine (HPG) to allow labelling of nascent polypeptides. Cells were then fixed, and Alexa Fluor was attached to HPG by Click chemistry. Samples were counter-stained with Hoechst and CellTracker Orange CMTMR, imaged, and analysed as described previously.

### Translation assay *in vitro*

*In vitro* translation assay was done with the Rabbit Reticulocyte Lysate (Promega, L4960) according to the manufacturer’s protocol. In short, translation reaction components were mixed with luciferase mRNA, and DMSO, SALSa or cycloheximide. The reactions were incubated 90 minutes at 30°C, followed by the addition of luciferin, a luciferase substrate. Luminescence was read using Infinite M200PRO plate reader (Tecan) until the signal stabilised.

### Ribosome profiling

Exponentially growing U2OS cells were treated with 10 *µ*M of SALSa for 48 h. Ribosomes were stalled by addition of 100 μg/mL cycloheximide for 5 min, and cells lysed in polysome lysis buffer (15 mM Tris-HCl pH 7.4, 15 mM MgCl_2_, 300 mM NaCl, 1% Triton-X-100, 0.1% β-mercaptoethanol, 200 U/mL RNAsin (Promega), 1 complete Mini Protease Inhibitor Tablet (Roche) per 10 mL). Nuclei were removed by centrifugation (9300 x g, 4°C, 10 min) and the cytoplasmic lysate was loaded onto a sucrose density gradient (17.5–50% in 15 mM Tris-HCl pH 7.4, 15 mM MgCl_2_, 300 mM NaCl and, for fractionation from BMDM, 200 U/mL Recombinant RNAsin Ribonuclease Inhibitor (Promega)). After ultracentrifugation (2.5 h, 35 000 rpm at 4°C in a SW60Ti rotor), gradients were eluted with a Teledyne Isco Foxy Jr. system into 13 fractions of similar volume.

### Transmission Electron Microscopy

Cells were fixed with 3% glutaraldehyde in 0.12 M phosphate buffer (PB) for 30 min at room temperature, scraped and collected in Eppendorf tubes, centrifuged to obtain a pellet, washed in PB, and postfixed in 2% osmium tetroxide. Cell samples were then dehydrated with acetone and embedded in ACM Durcupan (Fluka, Sigma-Aldrich) resin. Ultrathin sections (50–70 nm) were obtained using the UltraCut UC7 ultramicrotome (Leica, Microsystems, Germany), picked up on copper grids, stained with uranyl acetate and lead citrate, and examined with the JEM 1011 (JEOL, Japan) electron microscope operating at 80 kV. Micrographs were taken with a camera (Orius 1200A; Gatan, USA) using the DigitalMicrograph software package (Gatan, USA). Electron micrographs were processed using Adobe Photoshop CS6 (v.13.0.1) (Adobe Systems).

### Ribophagy assay in U2OS RPS3^Keima^ cell line

U2OS cells expressing RPS3^Keima^ protein were treated for 48 hours with 250 nM Torin1 or 10 *µ*M SALSa. Following the treatment, cells were stained with 2.5 *µ*g/ml Hoechst 3342 (Thermo Scientific, 62249) for 10 minutes. The medium was replaced and cells were imaged using Opera Phenix microscope (PerkinElmer) to capture Hoechst (Ex_395_/Em_480_), neutral-pH Keima (Ex_442_/Em_620_), and acidic-pH Keima (Ex_561_/Em_620_) signals. Images were analysed using Harmony software (PerkinElmer).

### Sequential HaloTag-based labelling of RPL29

Sequential HaloTag-based labelling of RPL29 was carried out as previously described^20^. Briefly, HCT116-RPL29^HaloTag^ cells were incubated for 1 hour with 100 mM HaloTag TMR ligand (Promega), followed by 3 PBS washes, and 24-hour incubation with 50 nM HaloTag R110 ligand (Promega) in the presence of DMSO or 10 *µ*M SALSa. The cells were then fixed with 4% formaldehyde and imaged with Opera Phenix microscope (PerkinElmer). Images were analysed using Harmony software (PerkinElmer).

### Visualisation of SALSa in cells

U2OS cells were treated for 2 hours with 10 *µ*M SALSa functionalised with alkyne moiety. To visualise the compound, fixed and permeabilised cells were incubated with click chemistry labelling solution (1 mM CuSO_4_, 50 *µ*M azide-AlexaFluor^594^, 5 mM sodium ascorbate; in PBS) for 1 hour, followed by PBS wash. Afterwards, cells were blocked, immunostained for Nucleophosmin or LAMP1, and counter stained with Hoechst 3342 as described above.

### RT-qPCR

Total RNA was extracted from cell pellets using a Purelink RNA Mini Kit (Invitrogen, no. 2183025) following the manufacturer’s instructions. Reverse-transcription and PCR amplification were performed using a TaqMan RNA-to-CT 1-Step Kit and the StepOnePlus Real-Time PCR Instrument (Applied Biosystems, Fisher Scientific). All primers used in this study are listed in **Table S1**.

### Northern blotting (agarose, PA)

Total RNA was extracted from drug-treated U2OS cells using TRIzol reagent (Invitrogen, 15596018) according to manufacturer’s protocol. RNA samples were heated at 80°C for 5 minutes in 6x TriTrack loading dye (Thermo Scientific, R1161) supplemented with GelRed (VWR, 41003) and stored on ice. Samples (5 *µ*g) were resolved either on a 1.2% agarose-TBE or a 4-20% polyacrylamide-TBE gel (Invitrogen, EC6225BOX). Agarose gels were electophoresed at 90 V for 120 min in 1x TBE, followed by a 30-min transfer onto Nylon Hybond membranes (Roche, 112092999001) using Trans-Blot SD Semi-Dry Electrophoretic Transfer cell (Bio-Rad) at 1.5 mA/cm^2^ at 4°C. Polyacrylamide gels were electrophoresed at 140 V for 90 min in 1x TBE, followed by a 2-hour wet transfer at 30 V. Membranes were rinsed with 2X SSC buffer (300 mM NaCl, 30 mM Na-citrate), crosslinked with UV light and incubated for 2 hours at 42°C with pre-hybridization solution (1x SSPE (150 mM NaCl, 10 mM Na_2_HPO_4_, 1 mM EDTA), 1x Denhardt’s solution (0.02% polyvinylpyrrolidone, 0.02% Ficoll 400, 0.02% BSA), 1% SDS, 50 *µ*g/ml salmon sperm DNA). Afterwards, 5’-biotinylated rRNA-specific probes (50 nM) were hybridized overnight at 49°C in hybridization solution (1x SSPE, 1% SDS). Membranes were then washed 3 x 15’ with low stringency buffer (6x SSPE, 0.2% SDS) and 3 x 15’ with high stringency buffer (1x SSPE, 0.2% SDS); all washes were done at 49°C. Signal was developed using Chemiluminescent Nucleic Acid Detection kit (Thermo Scientific, 89880) according to manufacturer’s protocol and imaged in Amersham Imager 600 (GE Healthcare). All hybridization probes are listed in **Table S1**.

### Circularised RNA RT-PCR

Total RNA was extracted from drug-treated U2OS cells using TRIzol reagent (Invitrogen, 15596018) according to manufacturer’s protocol. Circularisation was done using T4 RNA Ligase (NEB, M0204S) in 20 *µ*l reactions and 5 *µ*g input RNA at 37°C for 1 hour. RNA was purified by extraction with phenol:isoamyl alcohol:chloroform and precipitation with isopropanol. Next, reverse transcription was done using Super Script IV Reverse Transcriptase (Invitrogen, 18090050) according to manufacturer’s protocol with 250 ng inpur circularised RNA and 5.8S rRNA primer. Finally, the cDNA was used for end-point PCR to analyse regions flanking 5.8S rRNA. All primers used in this study are listed in **Table S1**.

### SALSa chemical affinity proteomics

SALSa^alk^ was conjugated to biotin by click reaction containing (per 1 pull-down sample) 5 pmol SALSa[alkyne], 5 pmol azide-PEG3-biotin (Jena Bioscience, CLK-AZ104P4-25), 10 mM CuSO_4_, 25 mM THPTA and 50 mM sodium ascorbate). After 2 h incubation at 45°C, 25 *µ*l pre-washed Dynabeads MyOne Streptavidin C1 were added, the sample was incubated for 2 hours at 25°C, and the beads were washed twice with PBS and twice with Pierce IP Lysis Buffer (Thermo Scientific, 87787). U2OS cells (20 × 10^6^ cells per sample) were suspended in Pierce IP Lysis Buffer supplemented with Halt Protease Inhibitor Cocktail (Thermo Scientific, 78430) and 0.1% Benzonase (Merck, E1014) and sonicated on ice at low intensity. Lysates were centrifuged and concentration measured using the DC Protein Assay Kit II (Bio-Rad, 5000112). Five mg of lysate was mixed with 25 *µ*l mock or SALSa^alk^-conjugated beads and incubated for 1 hour at 4°C with rotation. Beads were washed thrice with 1 ml IP Lysis Buffer and stored at −80°C until analysis. Proteins were reduced and alkylated by adding 50 *µ*l of 6M urea, 50 mM TEAB pH 8.5, 15 mM TCEP, 50 mM CAA to the dry beads and incubating them for 1 h in the dark at RT. Proteins were digested by adding 200ng of Lys-C (Wako) per sample for 4h at RT (estimated ratio enzyme:protein 1:50), followed by 200ng of trypsin (Promega) (estimated ratio enzyme:protein 1:50) incubated for 14 additional hours at 37 °C (1M urea, 50 mM TEAB pH 8.5). Resulting peptides were desalted using C18 stage-tips, speed-vac dried and re-dissolved in 21 *µ*l of 0.5% formic acid.

LC-MS/MS was done by coupling an UltiMate 3000 *RSLCnano* LC system to a Q Exactive Plus mass spectrometer (Thermo Fisher Scientific). Peptides were loaded into a trap column (Acclaim™ PepMap™ 100 C18 LC Columns 5 *µ*m, 20 mm length) for 3 min at a flow rate of 10 *µ*l/min in 0.1% FA. Then, peptides were transferred to an EASY-Spray PepMap RSLC C18 column (Thermo) (2 *µ*m, 75 *µ*m x 50 cm) operated at 45 °C and separated using a 60 min effective gradient (buffer A: 0.1% FA; buffer B: 100% ACN, 0.1% FA) at a flow rate of 250 nL/min. The gradient used was, from 2% to 6% of buffer B in 2 min, from 6% to 33% B in 58 minutes, from 33% to 45% in 2 minutes, plus 10 additional minutes at 98% B. Peptides were sprayed at 1.5 kV into the mass spectrometer via the EASY-Spray source and the capillary temperature was set to 300 °C. The mass spectrometer was operated in a data-dependent mode, with an automatic switch between MS and MS/MS scans using a top 15 method. (Intensity threshold ≥ 4.5e4, dynamic exclusion of 25 sec and excluding charges unassigned, +1 and > +6). MS spectra were acquired from 350 to 1500 m/z with a resolution of 70,000 FWHM (200 m/z). Ion peptides were isolated using a 2.0 Th window and fragmented using higher-energy collisional dissociation (HCD) with a normalized collision energy of 27. MS/MS spectra resolution was set to 35,000 (200 m/z). The ion target values were 3e6 for MS (maximum IT of 25 ms) and 1e5 for MS/MS (maximum IT of 110 msec).

Raw files were processed with MaxQuant (v 1.6.10.43) using the standard settings against a human protein database (UniProtKB/Swiss-Prot, 20,373 sequences). Carbamidomethylation of cysteines was set as a fixed modification whereas oxidation of methionines and protein N-term acetylation were set as variable modifications. Minimal peptide length was set to 7 amino acids and a maximum of two tryptic missed-cleavages were allowed. Results were filtered at 0.01 FDR (peptide and protein level). Afterwards, the “proteinGroups.txt” file was loaded in Prostar ^54^ using the LFQ intensity values for further statistical analysis. Briefly, proteins with less than 66% valid values in at least one experimental condition were filtered out. Missing values were imputed using the algorithms SLSA for partially observed values and DetQuantile for values missing on an entire condition. Differential analysis was performed using the empirical Bayes statistics Limma. Proteins with a p.value < 0.01 and a log2 ratio >1 or <-1 were defined as regulated. The FDR was estimated to be below 10.7%.

### Differentiation of SH SY5Y^PR97^

SY-SY5Y^PR97^ cells were seeded onto gelatine-coated 96-well plates (Greiner Bio-One, 655090) in Neurobasal™ (Gibco, 21103049), supplemented with 10% tetracycline-free FBS (PAN Biotech, P30-3602), 1% penicillin-streptomycin, 1% L-glutamine, and 1x B27 Supplement (Gibco, 17504044). Differentiation was carried out for 4 days without further media change. To induce the expression of PR peptide, 20 ng/ml Doxycycline was added. SALSa (or vehicle) was added together with doxycycline. For microscopy, cells were fixed with 4% paraformaldehyde for 10 minutes, permeabilized with 0.5% Triton X-100 in PBS for 15 minutes and blocked with 2.5% BSA in PBS for 30 minutes. Primary antibody incubation was performed overnight at 4 °C using anti-βIII-tubulin (BioLegend, Cat# 801202, 1:2000). After PBS washes, cells were incubated for 3 hours at room temperature with Alexa Fluor 488-conjugated goat anti-mouse IgG (Invitrogen, A11001, 1:500) and DAPI. Images were acquired using an ImageXpress Pico Automated Cell Imaging System (Molecular Devices) with a 10X objective.

### Drosophila work

All *Drosophila melanogaster* strains were obtained from the Bloomington Drosophila Stock Center (BDSC, Indiana, USA). The line used to model *C9orf72*-related Amyotrophic Lateral Sclerosis (ALS) and Frontotemporal Dementia (FTD) was BDSC #84727, carrying the construct P[UAS-(GGGGCC)49]48-10, which expresses 49 pure GGGGCC repeats under the control of the UAS promoter. This number of repeats has been reported to drive neurodegeneration in flies and to recapitulate the pathogenic G4C2 repeat expansion found in human *C9orf72*-associated ALS/FTD^55^. For neuronal expression, this strain was crossed with the pan-neuronal driver ELAV-GAL4 (BDSC #458), resulting in the experimental genotype UAS-(GGGGCC)49/ELAV-GAL4. As a control, ELAV-GAL4 flies were crossed with a strain carrying a pVal20 vector to knock down GAL4 (BDSC #35784), resulting in the Control (UAS-iGal4/ELAV-GAL4) genotype. For muscle-specific expression, the P[UAS-(GGGGCC)49]48-10 strain was crossed with the muscle driver MEF2, MEF2-GAL4 (BDSC #27390), resulting in the experimental genotype UAS-(GGGGCC)/MEF2-GAL4. As a control, MEF2-GAL4 flies were crossed with a strain carrying a pVal20 vector to knock down GAL4 (BDSC #35784), resulting in the Control (UAS-iGal4/MEF2-GAL4) genotype. Flies were maintained at 22°C, 70% humidity, under a 12 h light/dark cycle. Experimental flies were reared at 26 °C under the same humidity and light conditions.

SALSa treatment was done by incorporation of the compound into standard *Drosophila* food. Treatments started at the mating day and continued throughout adulthood. Food vials were replaced every 3–4 days to ensure compound stability and treatment effectivity. For lifespan analysis, 50 adult female flies per genotype were collected within 24 h of eclosion and maintained in groups of five per tube. Dead flies were counted every two days, and fresh food was provided weekly. Survival curves were generated using the Mantel–Cox (log-rank) test.

## Data availability

Mass spectrometry data have been deposited to the ProteomeXchange Consortium via the PRIDE partner repository^56^ with the dataset identifiers PXD075287 (for the chemical affinity purification) and PXD075027 (for PISA data). RNAseq data have been deposited at NCBI GEO (GSE324761).

## Acknowledgments

The authors want to thank the Chemical Biology Consortium Sweden (Karolinska Institutet) for providing screening library and chemistry support, the Chemical Proteomics unit (Karolinska Institutet) for PISA proteomics, Dr Gorka Gereñu (Miaker Developments) for Drosophila work, Ivó Hernández for help with RNAseq analyses and the proteomics, genomics and confocal microscopy units of the CNIO for their technical support. Work done in the O.F-C. lab was supported by grants from the Swedish Research Council (VR) (538-2014-31), the Spanish Ministry of Science, Innovation and Universities (PID2024-163103OB-I00, co-financed with European FEDER funds), La Caixa Foundation (HR22-00890), CIBERNED (CB24/05/00022) and a Karolinska Institutet Research Grant (awarded to B.P.).

## Author Contributions

B.P. performed most of the experiments and data analysis, prepared figures and contributed to the experimental plan, supervision of the study and writing of the manuscript; M. H., C.H. and J.C.-P. performed the chemical screens and initial analyses; V.L. performed the Keima, HaloTag, proteostat and polysome fractionation experiments; I. V. helped with polysome analyses; L.L. and S.R. provided technical help; Martin Haraldsson provided chemistry support; A.S. and S.M. performed Drosophila experiments; A.S. performed the SH-SY5Y^PR97^ experiments; M.L. helped with TEM analysis; D.H. helped to coordinate the study; O.F. conceived, coordinated and supervised the study, and wrote the manuscript.

