## Supplementary material for "A chemical inducer of ribophagy limits the toxicity of ALS-related arginine-rich peptides": Figures S1-S5

Porebski *et al* **Fig S1**

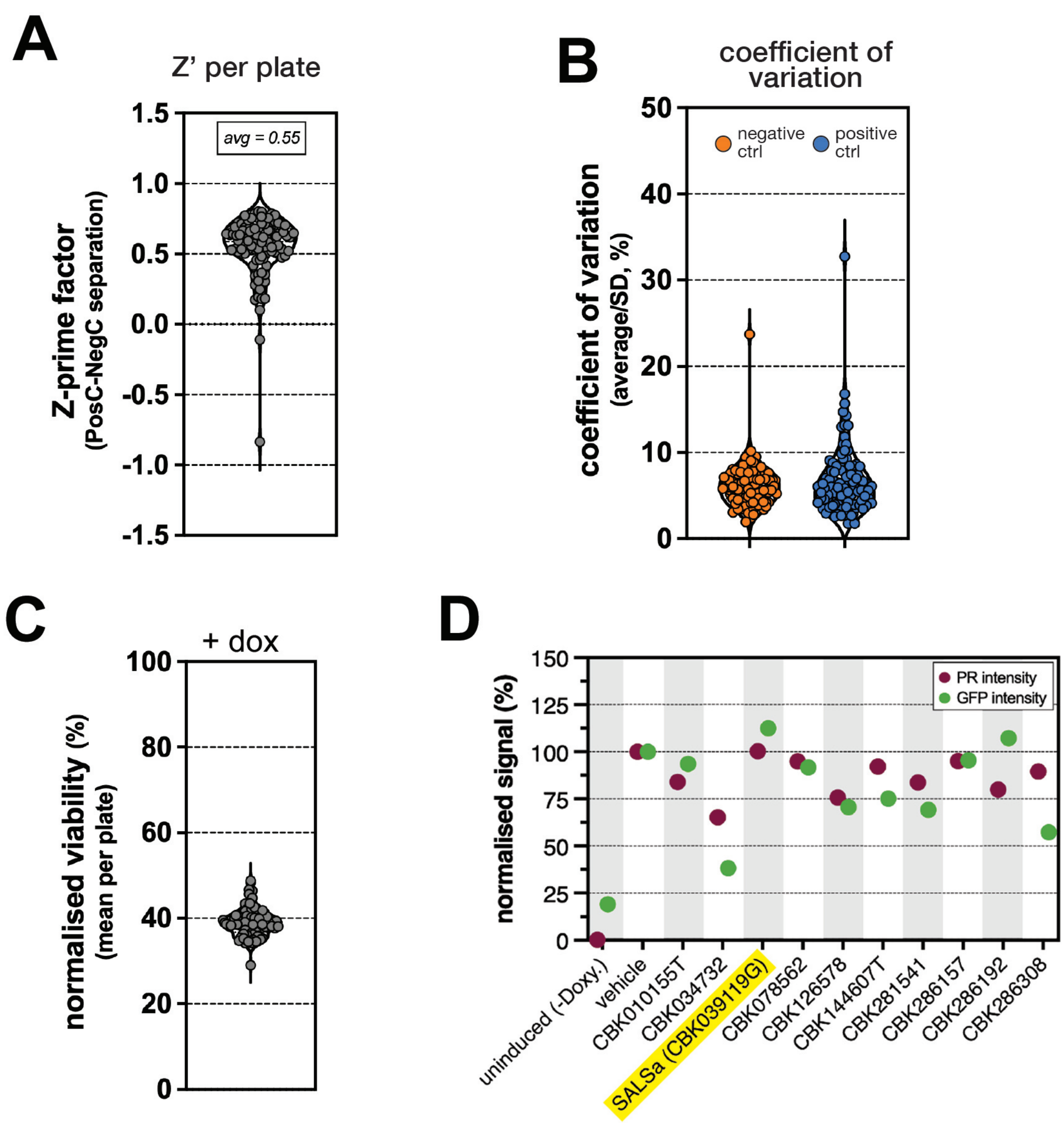

**Figure S1. Quality controls and validations of the chemical screen.**

A. Violin plot showing the distribution of the Z' factor across the screen. Each dot represents one screening plate. Z' values were calculated based on 16 wells of positive and negative controls.

B. Violin plot showing the distribution of the coefficient of variation (CV) across the screen for both positive and negative controls. Each dot represents one screening plate. CV values were calculated based on 16 well of positive and negative controls.

C. Violin plot showing the distribution of the dox-induced decrease in the viability of $U2OS^{PR97}$ cells across the screen. Each dot represents one screening plate. Within each plate, mean of 16 negative control wells was normalised to the mean of 16 positive control wells.

D. Effect of 10 hit compounds (10 $\mu$M) on dox-inducible expression of $(PR)_{97}$ peptides (pink dots) and EGFP (green dots) in $U2OS^{PR97}$ and $U2OS^{EGFP}$ cells. Each dot represents mean intensity normalised to the signal from vehicle-treated induced sample.

Porebski *et al* **Fig S2**

**A**

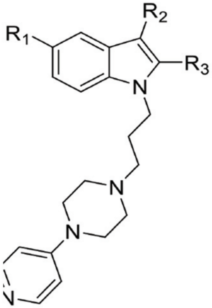


| # | $R_1$ | $R_2$ | $R_3$ | $EC_{50}$ (µM) |
|---|---|---|---|---|
| **1** | F | H | Me | 3.97 |
| **2** | H | H | H | 3.38 |
| **3** | F | H | H | 4.44 |
| **4** | Br | H | H | 1.35 |
| **5** | I | H | H | 1.05 |
| **6** | 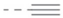 | H | H | 2.91 |
| **7** | H | 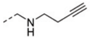 | H | 0.98 |
| **8** | H | CHO | H | 14.36 |
| **9** | H | H | 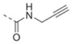 | inact. |

**B**

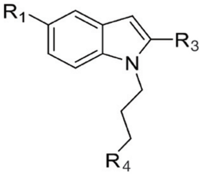


| # | $R_1$ | $R_3$ | $R_4$ | aliphatic $pKa_1$ | aryl ring $pKa_2$ | $EC_{50}$ (µM) |
|---|---|---|---|---|---|---|
| **1** | F | Me | 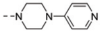 | 8.1 | 8.8 | 3.97 |
| **10** | H | H | 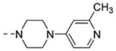 | 8.3 | 9.2 | 6.64 |
| **11** | Br | H | 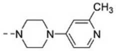 | 8.3 | 9.2 | 1.23 |
| **12** | F | Me | 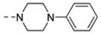 | 8.6 | - | inact. |
| **13** | F | Me | 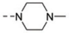 | 8.5 | - | inact. |
| **14** | F | Me | 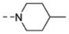 | 10 | - | inact. |
| **15** | F | Me | 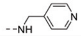 | 9.6 | 2.7 | inact. |
| **16** | F | Me | 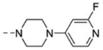 | 8.2 | 2.3 | inact. |
| **17** | F | Me | 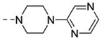 | 9.2 | 0.3 | inact. |
| **18** | F | Me | 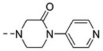 | 6.4 | 4.2 | inact. |
| **19** | H | H | 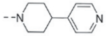 | 9.6 | 5.4 | inact. |

**C**

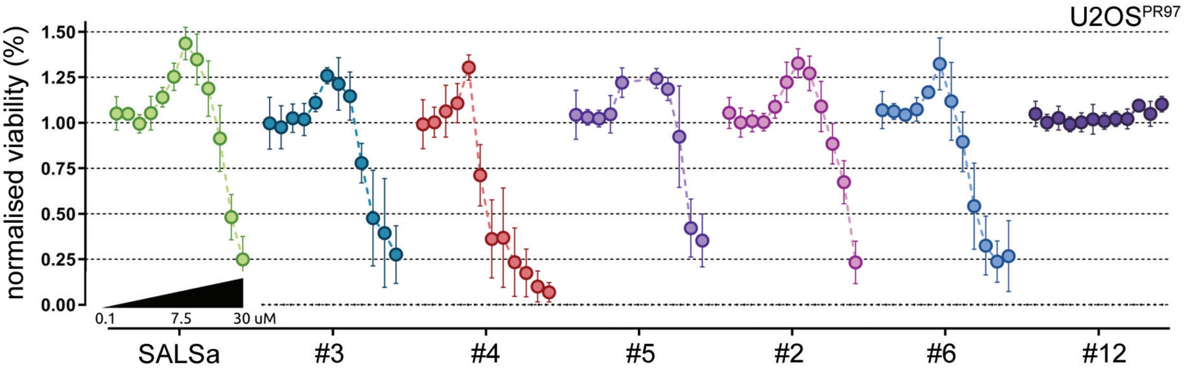

**Figure S2. Structure-activity relationship (SAR) analysis of SALSa.**

A. SAR analysis of the core indole ring. Positions R1 and R2 were tolerant to removal or small substitutions (2-8), while changes in position R3 had negative effect on compound's activity. Calculation of the half maximal effective concentration ($EC_{50}$) was based on the ascending phase of the dose-response curve, representing the benefit in survival.

B. SAR analysis of the two-ring system. Removal of basic amines or truncations rendered the compound inactive (12-14). Reinstallation of nitrogen in aryl ring with concomitant decrease in basicity did not recover the activity (15-19). Calculation of the half maximal effective concentration ($EC_{50}$) was based on the ascending phase of the dose-response curve.

A. Dose-response curves of SALSa and selected analogues rescue activity in $U2OS^{PR97}$ cells. Plotted is the viability normalised against dox-treated cells (N = 4).

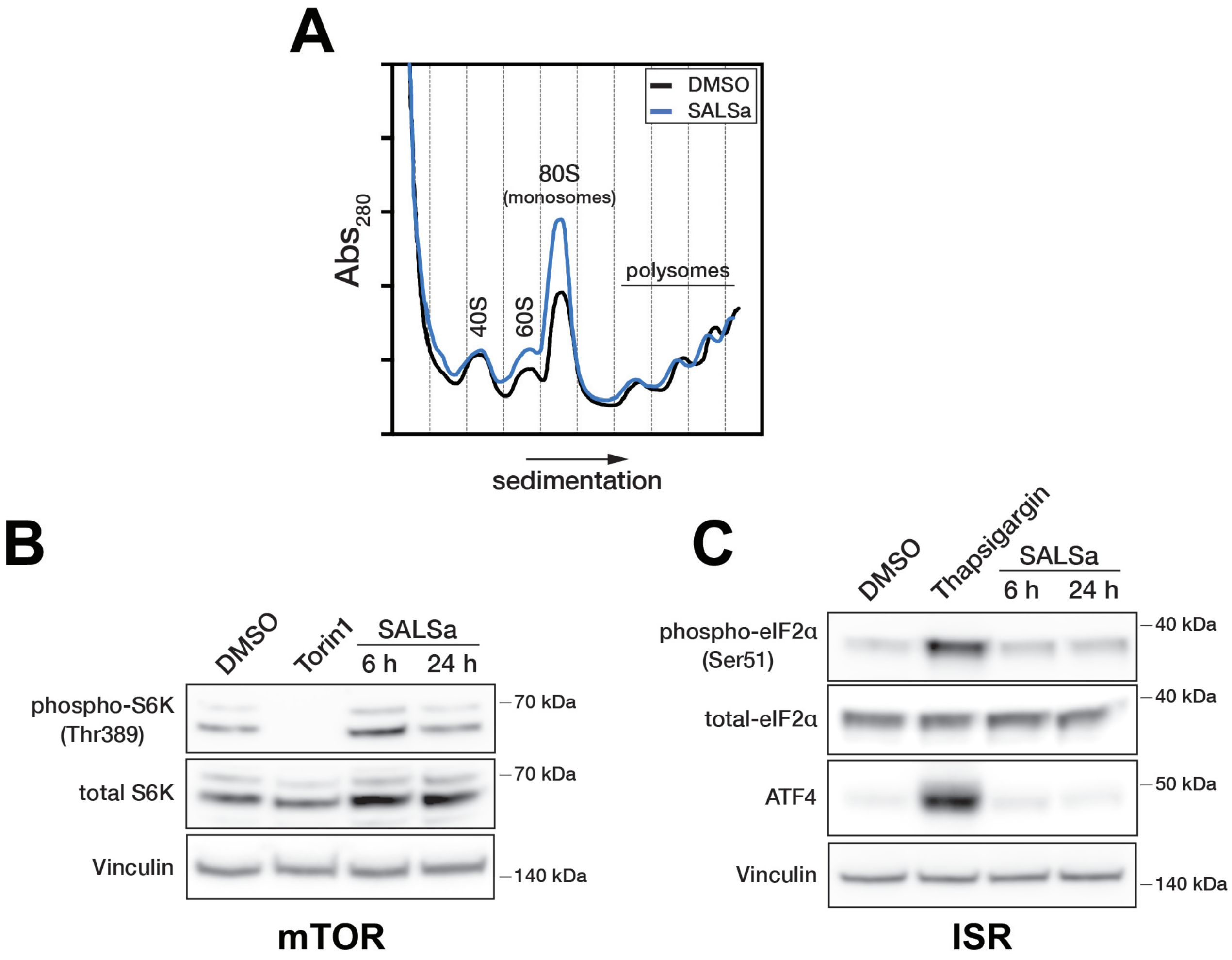
A
DMSO
SALSa
80S
(monosomes)
polysomes
40S
60S
Abs280
sedimentation
B
DMSO
Torin1
SALSa
6 h
24 h
phospho-S6K
(Thr389)
total S6K
Vinculin
70 kDa
70 kDa
140 kDa
mTOR
C
DMSO
Thapsigargin
SALSa
6 h
24 h
phospho-eIF2α
(Ser51)
total-eIF2α
ATF4
Vinculin
40 kDa
40 kDa
50 kDa
140 kDa
ISR

**Figure S3. Impact of SALSa on polysome assembly and mTOR/ISR signalling**

A. Polysome profiling of U2OS cells treated with DMSO or 10 $\mu$M SALSa for 24 h.
B. Western blotting analysis of mTOR signalling (S6 kinase phosphorylation) in U2OS cells treated with DMSO, 100 nM Torin1 or 10 $\mu$M SALSa for 24h. Vinculin levels are shown as a loading control.
C. Western blotting analysis of (ISR) signalling (eIF2$\alpha$ phosphorylation and ATF4 levels) in U2OS cells treated with DMSO, 2 $\mu$M Thapsigargin or 10 $\mu$M SALSa for 24h. Vinculin levels are shown as a loading control.

Porebski *et al* **Fig S4**

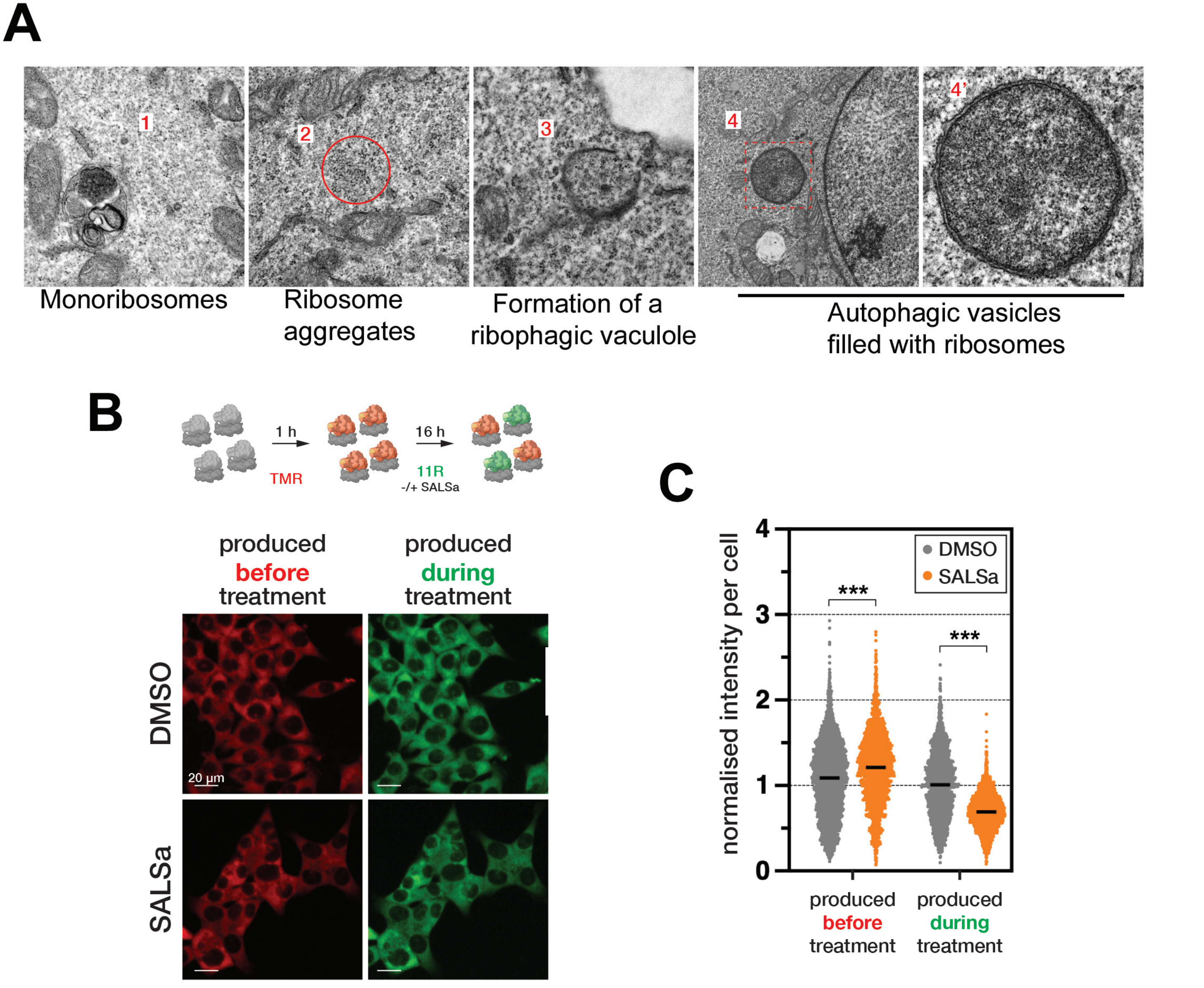

**Figure S4. SALSa induces ribosomal aggregation and the autophagic clearance of ribosomes.**

A. Transmission electron microscopy images of U2OS cells treated with 10 $\mu$M SALSa for 4h. Various stages that go from the formation of ribosomal aggregates to the formation of ribophagy vesicles are shown.

A. Microscopy images of HCT116-RPL29$^{HaloTag}$ cells. The cells were first labelled with TMR fluorophore for 1h to mark existing ribosomes, followed by a 16h labelling with 11R fluorophore (to tag newly generated ribosomes) in the presence or absence of 10 $\mu$M SALSa.

B. Quantification of sequentially labelled RPL29$^{HaloTag}$ proteins. For each cell, the signal intensity representing pre-existing or nascent ribosomes is plotted. Statistical significance was determined by two-way ANOVA. ***, $p < 0.001$,

Porebski *et al* **Fig S5**

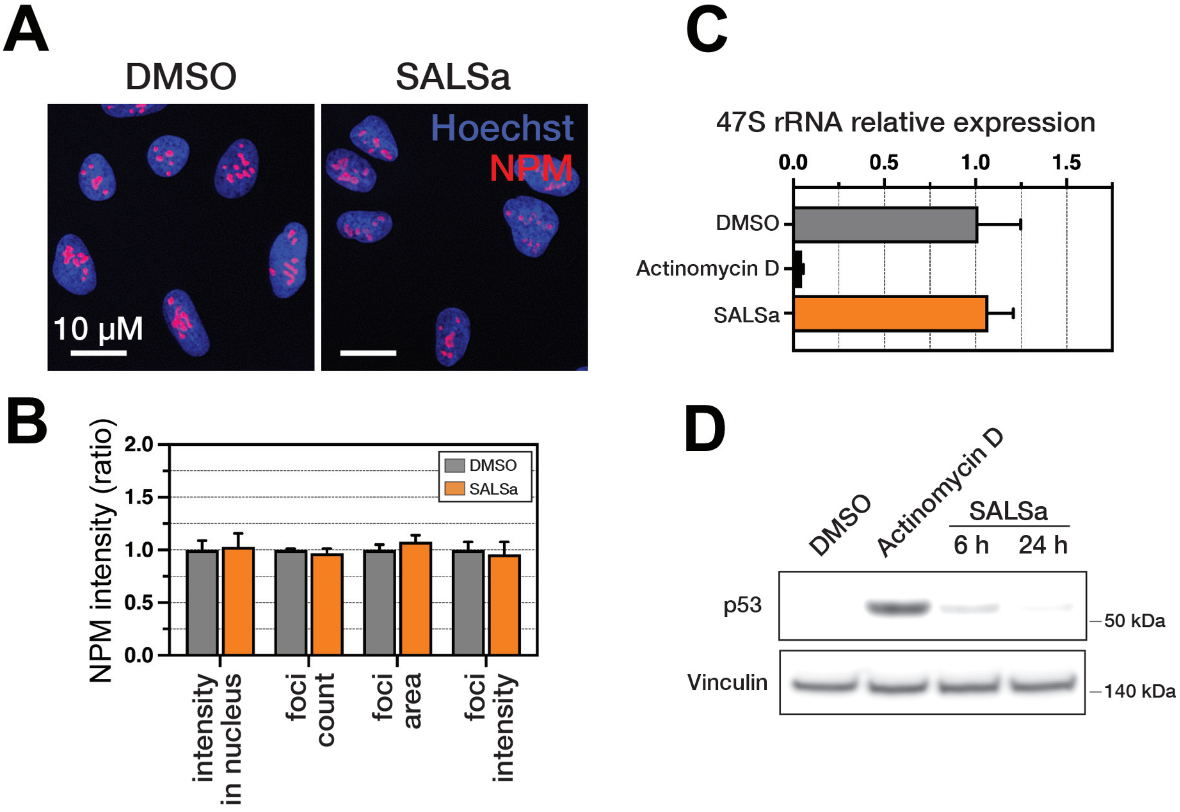

**Figure S5. SALSa does not affect nucleolar integrity**

A. Microscopy images of U2OS cells treated with 10 $\mu$M SALSa for 6 h. Cells were stained with Hoechst 3342 (blue) to visualise nuclei and anti-Nucleophosmin (NPM) to stain nucleoli (red). Scale bar 10 $\mu$m.

B. Quantification of various metrics of nucleolar morphology based on the NPM staining, in U2OS cells treated with 10 $\mu$M SALSa for 6 h. Plotted values represent the mean of 3 replicates normalised to the DMSO sample. For each replicate at least 150 cells were analysed.

C. RT-qPCR analysis of 47S pre-rRNA levels in U2OS cells treated with DMSO, 5 nM Actinomycin D or 10 $\mu$M SALSa for 6 h. Plotted values represent mean of 3 replicates normalised to the DMSO sample.

D. Western blotting of P53 levels as a proxy for the nucleolar stress response in U2OS cells treated with DMSO, 5 nM Actinomycin D (6 h) or 10 $\mu$M SALSa. Vinculin levels are shown as a loading control.
