## Supplementary material for "A chemical inducer of ribophagy limits the toxicity of ALS-related arginine-rich peptides": Table S1

**TABLE S1.** LIST OF REAGENTS USED IN THIS STUDY.

| REAGENT or RESOURCE | SOURCE | IDENTIFIER |
|---|---|---|
| **Antibodies** | | |
| anti-ATF4 | CST | 11815 |
| anti-eIF2α | CST | 9722 |
| anti-eIF2α-phospho (S51) | CST | 9721 |
| anti-LAMP1 | CST | 15665 |
| anti-LC3 | Sigma | L7543 |
| anti-NPM | Abcam | ab10530 |
| anti-p53 | Abcam | ab1101 |
| anti-Pro-Arg | Proteintech | 23979-1-AP |
| anti-RPL11 | Thermo Scientific | 37-3000 |
| anti-RPL11 | Proteintech | 14583-1-AP |
| anti-RPL37A | Atlas Antibodies | HPA065327 |
| Anti-RPS6 | CST | 2217 |
| anti-S6K | CST | 9202 |
| anti-S6K-phospho (T389) | CST | 9205 |
| anti-TUBB3 | BioLegend | 801202 |
| anti-Vinculin | Abcam | ab129002 |
| anti-Mouse-Alexa488 | Invitrogen | A11001 |
| anti-Rabbit-Alexa647 | Invitrogen | A-21244 |
| anti-Rabbit-HRP | Invitrogen | 11859140 |
| **Chemicals, peptides, and recombinant proteins** | | |
| Actinomycin D | Sigma | A1410 |
| Azide-PEG3-biotin | Jena Bioscience | CLK-AZ104P4-25 |
| B27 supplement | Gibco | 17504044 |
| Blasticidin | InvivoGen | ant-bl-05 |
| CellTiter-Glo | Promega | G75751 |
| CellTracker Orange CMTMR | Thermo Scientific | C2927 |
| Cycloheximide | Sigma | C4859 |
| DAPI | Invitrogen | D1306 |
| Doxycycline | Sigma | D9891 |
| HaloTag R110 | Promega | G3221 |
| HaloTag TMR | Promega | G8251 |
| Hoechst | Thermo Scientific | 62249 |
| Methionine-free DMEM | Gibco | 21013024 |
| Proteostat | Enzo | ENZ-51023 |

**TABLE S1.** LIST OF REAGENTS USED IN THIS STUDY.

| | | |
|---|---|---|
| Retinoic acid | Sigma | R2625 |
| Super Script IV Reverse Transcriptase | Invitrogen | 18090050 |
| T4 RNA Ligase | NEB | M0204S |
| Tetracycline-free FBS | Takara Bio | 631106 |
| Tetracycline-free FBS | PAN Biotech | P30-3602 |
| Thapsigargin | Abcam | ab120286 |
| THPTA | Jena Bioscience | CLK-1010-1G |
| Torin1 | Tocris | 4247 |
| TRIzol reagent | Invitrogen | 15596018 |
| **Critical commercial assays** | | |
| Chemiluminescent Nucleic Acid Detection kit | Thermo Scientific | 89880 |
| Click-iT HPG Alexa Fluore Protein Synthesis Assay | Invitrogen | C10428 |
| POWER SYBR Green RNA-to-CT 1-Step kit | Thermo Scientific | 4389986 |
| PureLink RNA Mini kit | Invitrogen | 12183025 |
| QuantSeq 3' mRNA-Seq Library Prep Kit | Lexogen | N/A |
| Rabbit Reticylocyte Lysate | Promega | L4960 |
| **Deposited data** | | |
| Chemical affinity proteomics data | This study | PRIDE: PXD075287 |
| PISA proteomics data | This study | PRIDE: PXD075027 |
| RNA-Seq data | This study | NCBI GEO: GSE324761 |
| **Experimental model: Cell lines** | | |
| HCT116-RPL29-HaloTag9 | An et al. | |
| MCF-7 | ATCC | HTB-22 |
| SH SY5Y-PR97 (TetON | This study | |
| U2OS | ATCC | HTB-96 |
| U2OS-EGFP (TetON) | Colicchia et al. | |
| U2OS-PR97 (TetON) | Sirozh et al. | |
| U2OS-RPS3-Keima | This study | |
| **Experimental model: Organisms/strains** | | |
| Drosophila: ELAV-GAL4 | BDSC | $458; RRID:BDSC_458 |
| Drosophila: Mef2-GAL4 | BDSC | #27390; RRID:BDSC_27390 |
| Drosophila: P{UAS-(GGGGCC)49}48-10 | BDSC | #84727; RRID:BDSC_84727 |
| Drosophila: pVal20-GAL4-RNAi | BDSC | #35784; RRID:BDSC_35784 |
| Drosophila: UAS-(GGGGCC)49/ELAV-GAL4 | Miaker Developments | |

**TABLE S1.** LIST OF REAGENTS USED IN THIS STUDY.

| | | |
|---|---|---|
| Drosophila: UAS-(GGGGCC)49/Mef2-GAL4 | Miaker Developments | |
| **Oligonucleotides** | | |
| 47S qPCR fw: GAACGGTGGTGTGTCGTT | Kwon et al. | |
| 47S qPCR rev: GCGTCTCGTCTCGTCTCACT | Kwon et al. | |
| ActB qPCR fw: TCACAATGTGGCCGAGGACTTT | Espinoza et al. | |
| ActB qPCR rev: AGAAGTGGGGTGGCTTTTAGGA | Espinoza et al. | |
| ITS1 NB probe: [Btn]-GGCCTCGCCCTCCGGGCTCCGTTAATGAT | O'Donohue et al. | |
| ITS2 NB probe: [Btn]-CTGCGAGGGAACCCCCAGCCGCGCA | O'Donohue et al. | |
| 5.8S RT on circRNA: CGAACGCACTTGCGG | This study | |
| 5.8S fw: CGTCGCTTGCCGATC | This study | |
| 5.8S rev: GATGATCAATGTGTCCTGC | This study | |
| **Recombinant DNA** | | |
| pLENTI-RPS3-Keima | Addgene | 127140 |
| **Software and algorithms** | | |
| BioJupies | Torre et al. | https://maayanlab.cloud/biojupies/ |
| CellProfiler 4.0 | Stirling et al. | https://cellprofiler.org |
| GraphPad Prism | Version 11 | https://www.graphpad.com |
| KNIME Analytics Platform | Berthold et al. | https://www.knime.com |
| STRING | | https://www.string-db.org |
| **Other** | | |
| Dynabeads MyOne Streptavidin C1 | Invitrogen | 65-001 |
